# Harnessing anti-CRISPR to suppress, subtract, and segregate Cas9 activity for precision CRISPR in *Drosophila*

**DOI:** 10.64898/2025.12.07.692848

**Authors:** Yifan Shen, Michael Sheen, Ann T. Yeung, Xinchen Chen, Bei Wang, Zixian Huang, Claire A. Ho, Zachary I. Lakkis, Chun Han

**Affiliations:** Department of Molecular Biology and Genetics, Cornell University, Ithaca, New York, USA Weill Institute for Cell and Molecular Biology, Cornell University, Ithaca, New York, USA

**Keywords:** CRISPR/Cas9, Anti-CRISPR, *Drosophila* genetics, genome editing, tissue-specific CRISPR, germline, soma, balancer

## Abstract

Tissue-specific CRISPR (ts-CRISPR) is a powerful approach for studying cell and developmental biology by restricting mutagenesis to specific tissues. However, the precision of this approach is often compromised by non-specific, “leaky” Cas9 activity that confounds phenotypic analysis and destabilizes Cas9/gRNA stocks. To address these limitations in *Drosophila*, we developed a toolkit based on the Anti-CRISPR (Acr) protein AcrIIA4. We first identified AcrIIA4 as a potent *in vivo* Cas9 inhibitor with high stability and established the temporal requirements for its function. Based on these findings, we generated three classes of Acr tools. First, a collection of AcrIIA4-bearing balancers robustly suppresses Cas9 and enables the stable maintenance of complex Cas9/gRNA stocks. Importantly, maternal deposition of AcrIIA4 from these balancers provides a means of temporal control, delaying Cas9 activity until metamorphosis. Second, tissue-specific AcrIIA4 transgenes refine leaky Cas9 drivers in a “tissue-subtraction” strategy. Finally, germline-specific and soma-specific Acr tools efficiently segregate Cas9 activity, solving bidirectional leakiness between these two compartments. This comprehensive AcrIIA4 toolkit provides new levels of precision, versatility, and temporal control for *Drosophila* CRISPR applications.

## INTRODUCTION

The CRISPR/Cas9 system has revolutionized *Drosophila* genetics by enabling powerful and convenient genome editing across diverse experimental paradigms ^1^. Among the numerous applications, the most frequent use of CRISPR/Cas9 is to knock out a gene of interest. In this process, Cas9 generates double-strand breaks (DSBs) at the gene locus ^2^, and the DSBs are subsequently repaired by the error-prone non-homologous end joining (NHEJ) pathway to generate insertion-deletion (indel) mutations ^3, 4, 5^. Such an approach is very effective for generating inheritable mutations with germline-expressed Cas9 and guide RNAs (gRNAs) ^6, 7^. Importantly, CRISPR/Cas9 also enables tissue-specific gene knockout (KO) in whole organisms. In such applications, Cas9, or gRNAs, or both are expressed in a tissue-specific manner, resulting in biallelic loss-of-function (LOF) mutations restricted to the target tissue ^8, 9, 10^. Compared to the traditional analysis of whole-animal mutants, this tissue-specific CRISPR (ts-CRISPR) strategy avoids complications from early lethality, facilitates studies of cell autonomy and other complex phenotypes, and is conveniently implemented by simply combining Cas9 and gRNAs transgenes, regardless of the target gene’s location.

Despite the advantages of ts-CRISPR, its potential has not been fully realized due to critical limitations. Foremost, tissue-specific Cas9 lines, either driven by tissue-specific enhancers or expressed through the Gal4/UAS binary system, often exhibit problematic non-specific, or “leaky” activity in unintended tissues ^9, 11, 12, 13, 14^. This leakiness can undermine the specificity of phenotypic analysis, especially in developmental studies where a gene’s function may vary across different tissues and developmental stages. Similarly, Cas9 lines designed to be germline-specific often show secondary leaky activity in somatic tissues ^13^, which can cause unintended lethality and confound experiments requiring germline-specific genome editing. Furthermore, although maintaining Cas9 and gRNA transgenes in the same stock can greatly simplify downstream genetic crosses, this approach is associated with two significant risks: First, leaky Cas9 activity in the germline may mutate and permanently destroy the gRNA target sequence, leading to inactivation of CRISPR in future generations. Second, the Cas9/gRNA combination may result in sickness or lethality even with tissue-specific drivers (if the target gene is essential), making the stock difficult to maintain. Thus, tools that can precisely control Cas9 activity—removing it from unintended tissues and suppressing it entirely in stock animals—will be highly desirable.

Anti-CRISPR (Acr) proteins are a promising solution to these constraints. Acr proteins are small, diverse proteins encoded by phages and mobile genetic elements that can inhibit bacterial CRISPR-Cas immune systems, allowing the phages to evade host defense ^15^. Among the Acr families, the AcrIIA group is particularly notable because it targets type II-A CRISPR-Cas systems, such as *Streptococcus pyogenes* Cas9 (SpyCas9), the most widely used tool for genome editing in eukaryotes, including *Drosophila* ^1^. By expressing Acr transgenes in specific tissues, researchers can “subtract” leaky Cas9 activity and improve the tissue-specificity of ts-CRISPR. Furthermore, ubiquitously expressed Acr lines would allow Cas9 and gRNA transgenes to be co-maintained in stable, healthy stocks. Lastly, expressing Acr exclusively in germline or somatic tissues would enable precise segregation of Cas9 activity in these two compartments.

Although Acrs have been used in many cell culture studies ^16, 17, 18^, they have rarely been tested in whole animals ^19^. The *in vivo* properties of Acr proteins in *Drosophila* tissues thus remain largely uncharacterized. In this study, we systematically evaluated and developed Acrs into a robust and versatile toolkit to overcome these critical limitations in *Drosophila* ts-CRISPR. Through comparison of three Acr proteins in *Drosophila*, we identified AcrIIA4 as the most potent inhibitor and defined the temporal requirements for its effective suppression of Cas9. Based on these findings, we developed Acr-bearing balancers to stably maintain Cas9 and gRNA transgenes in the same line, to streamline ts-CRISPR experiments, and to utilize maternal AcrIIA4 deposition to delay tissue-specific knockout. Furthermore, we demonstrate that tissue-specific AcrIIA4 can be used in a tissue-subtraction strategy to enhance the tissue specificity of Cas9 lines. Finally, germline- and soma-specific AcrIIA4 lines enable separation of Cas9 activity between these two tissue types, facilitating more robust ts-CRISPR and a more efficient pipeline for Gal4-to-Cas9 conversion. Overall, these Acr tools improve the precision, versatility, and robustness of CRISPR applications in *Drosophila*. Similar strategies could be applied to other organisms.

## RESULTS

### AcrIIA4 can efficiently suppress cas9 activity in *Drosophila*

To identify an AcrII protein that can efficiently suppress Cas9 in *Drosophila*, we first optimized detection of Cas9 activity using a previously described reporter, *GSR* (for <u>G</u>FP <u>S</u>SA <u>r</u>eporter) (Fig. 1A) ^11^. This reporter features two incomplete EGFP repeats separated by a short sequence, which in turn is targeted by two gRNAs expressed by the same construct. Cas9 activity results in DSBs in the short sequence, which can then be repaired by single-strand annealing (SSA) between the EGFP repeats to reconstitute a complete EGFP coding sequence. The construct ubiquitously drives this cassette, which also includes a downstream nuclear GFP (nGFP) sequence linked by a 2A self-cleaving peptide, designed to mark the cytoplasm and nuclei of cells where Cas9 has been active ^11^. However, initial characterization of *GSR* using a ubiquitous *Actin-Cas9* driver ^6^ revealed incomplete labeling in larval tissues. For example, only 42% of the epidermal area in late 3^rd^ instar larvae was GFP positive (Fig. 1B, 1E). We reasoned that incomplete GSR activation could be due to the competing error-prone NHEJ repair pathway, which would create indel mutations at the target site rather than reconstituting the EGFP sequence. To test this, we introduced the reporter into an NHEJ-deficient *Lig4* (*DNA ligase IV*) mutant ^20^. Indeed, GFP labeling of the epidermis increased to 53% in *Lig4* heterozygous females and to 89% in hemizygous males (Fig. 1C-1E), revealing a strong, dosage-dependent competition from the NHEJ pathway that otherwise masks significant Cas9 activity detected by this SSA-based reporter. Based on these results, we decided to perform all subsequent somatic Cas9 activity assays with *GSR* in the *Lig4^-^/Y* background to maximize reporter sensitivity, unless otherwise noted.

**Figure 1.**
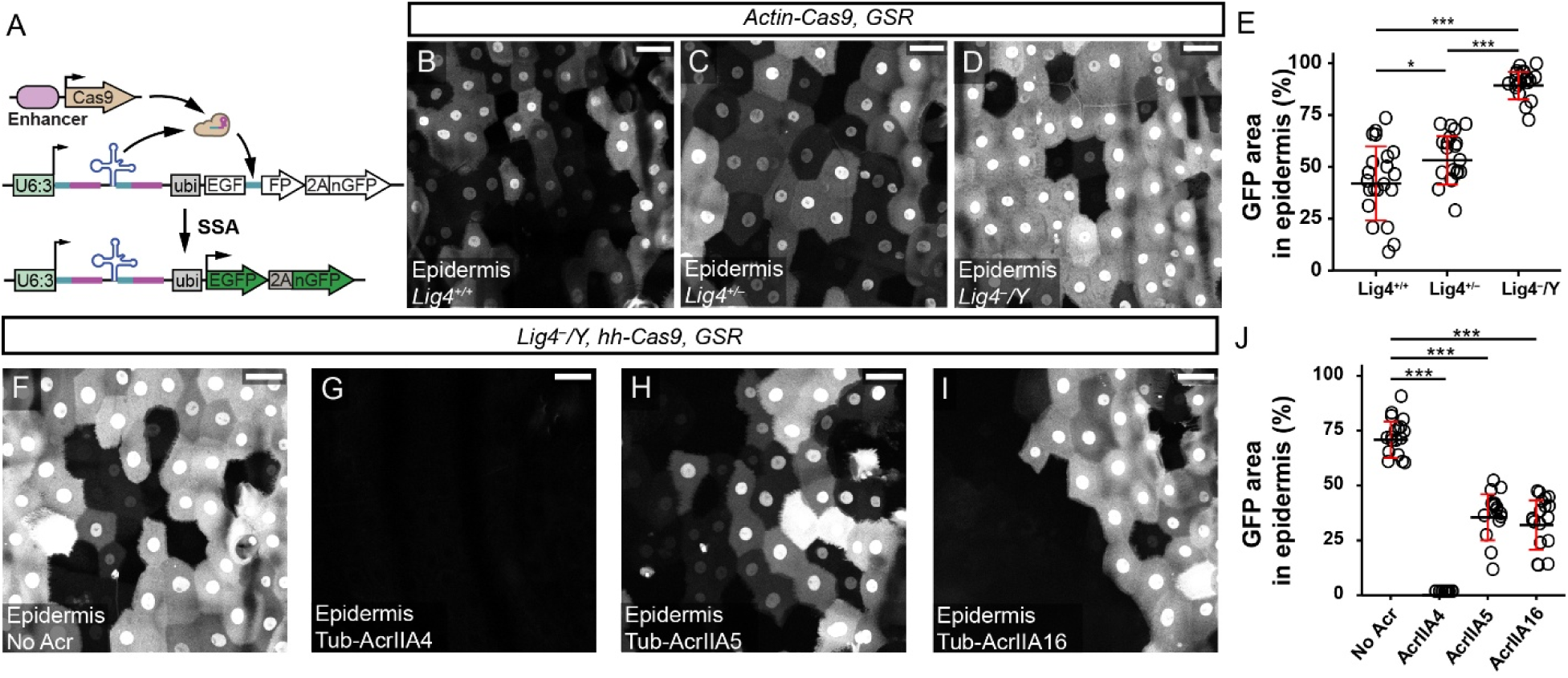
AcrIIA4 can efficiently suppress cas9 activity in *Drosophila*. (A) Design principle of *GSR*. SSA, single strand annealing. (B-D) *GSR* activation patterns by *Actin-Cas9* in the larval epidermis of a wildtype control (B), a heterozygote of *Lig4* null mutant (C), and a hemizygote of *Lig4* null mutant (D). (E) Percentage of the GFP-positive area in the larval epidermis of the indicated genotypes. n = number of larval segments: *Lig4^+/+^* (n = 21), *Lig4^+/-^* (n = 20), *Lig4^-^/Y* (n = 21). (F-I) *GSR* activation patterns by *hh-Cas9* in the epidermis without Acr (F), with *Tub-AcrIIA4* (G), with *Tub-AcrIIA5* (H), and with *Tub-AcrIIA16* (I). (J) Percentage of the GFP-positive area in imaged regions with different AcrIIA candidates. n = number of larval segments: No Acr (n = 18), AcrIIA4 (n = 18), AcrIIA5 (n = 16), AcrIIA16 (n = 18). In all plots, black bar, mean; red bar, SD. One-way analysis of variance (ANOVA) and Tukey’s honest significant difference (HSD) test. *p≤0.05, ***p≤0.001. Scale bar, 50 µm.

We selected three AcrII proteins previously known to suppress Cas9 activity, including AcrIIA4 ^21^, AcrIIA5 ^22^, and AcrIIA16 ^23^, and generated corresponding transgenes driven by a ubiquitous *Tub* promoter. We evaluated these transgenes using *hh-Cas9*, which is active in the posterior half of each segment in the larval epidermis ^8^. By itself, *hh-Cas9* activated *GSR* in 71% of the epidermal area in the examined regions (Fig. 1F, 1J). Co-expression of *Tub-Acr* transgenes reduced *hh-Cas9* activity to different extents: While AcrIIA5 and AcrIIA16 suppressed Cas9 activity to 36% and 32% areas, respectively, AcrIIA4 eliminated Cas9 labeling (Fig. 1F-1J). This demonstrates that AcrIIA4 is the most potent Cas9 inhibitor of the three tested in *Drosophila* somatic tissues. Based on these results, we selected AcrIIA4 for subsequent experiments and further characterization.

### AcrIIA4-bearing balancers efficiently suppress germline Cas9 activity

Many *Drosophila* genetic tools, including Cas9 lines, are maintained in stocks using balancers—chromosomes that suppress recombination ^24^. We reasoned that if these balancers also ubiquitously expressed AcrIIA4, they could be used to keep co-resident Cas9 and gRNA transgenes completely silent in stock animals. To that end, we generated *Tub-AcrIIA4* transgenes with fluorescent (*3xP3-DsRed*) or visible (*mini-white*) markers and hopped them onto commonly used balancers (Table 1) through P-element-mediated transposition.

**Table 1.** AcrIIA4-bearing balancers.

| ID | Full Genotype | Chr. | Marker |
| --- | --- | --- | --- |
| ACR1 | FM7a, P{RFP[DsRed.3xP3.cUa]=Tub-AcrIIA4.RFP+}17/C(1)DX, y[1] f[1] | X | 3XP3-DsRed |
| ACR2 | FM7a, P{RFP[DsRed.3xP3.cUa]=Tub-AcrIIA4.RFP+}46/C(1)DX, y[1] f[1] | X | 3XP3-DsRed |
| ACR3 | w[*]; sna[Sco]/CyO, P{RFP[DsRed.3xP3.cUa]=Tub-AcrIIA4.RFP+}27 | II | 3XP3-DsRed |
| ACR4 | w[*]; sna[Sco]/CyO, P{RFP[DsRed.3xP3.cUa]=Tub-AcrIIA4.RFP+}30 | II | 3XP3-DsRed |
| ACR5 | w[*]; TM3/TM6B, Tb[1], P{RFP[DsRed.3xP3.cUa]=Tub-AcrIIA4.RFP+}3r | III | 3XP3-DsRed |
| ACR6 | w[*]; wg[Sp-1]/CyO, P{Wee-P.ph0}2; TM2/TM6B, Tb[1] P{w[+mC]=Tub-AcrIIA4.w+}3w | III | Mini-white |

A primary application of such balancers is to prevent genetic instability of gRNA target sites. In stocks containing both Cas9 and gRNA transgenes, leaky Cas9 activity in the germline can mutate and permanently disrupt the gRNA target sequence, rendering the gRNA transgene ineffective in subsequent generations. An effective Acr balancer would prevent this outcome. To assess the efficacy of AcrIIA4 balancers in the germline, we devised a germline GSR assay: If a parent fly carries a germline-specific Cas9 and the GSR reporter, active Cas9 in the germline will induce SSA repair of the GSR allele, resulting in whole-body GFP-positive (GFP^WB^) progeny (Fig. 2A). An AcrIIA4 balancer added into the parent allows us to assay inhibition of germline Cas9 activity by quantifying the reduction in GFP^WB^ progeny. For this purpose, we first quantified the efficiency of GSR activation by two germline-specific Cas9s: *nos-Cas9* ^7^ and *bam-Cas9-P2A-FLP* (*bam-CF*) ^25^. *nos-Cas9* led to GSR activation in 89.5% of male and 95.3% of female germline transmissions, while *bam-CF* activated GSR in 93.2% of male and 93.9% of female transmissions (Fig. 2B and S1). The remaining progeny were either completely GFP-negative (GFP^-^) or showed sporadic, random GFP patterns (GFP^SR^) (Fig. S1A-S1D). Further analysis showed that GFP^-^ progeny carried either a mutated *GSR* allele (due to error-inducing NHEJ repair) or an intact *GSR* (due to precise repair in the parental germline) without the Cas9, while GFP^SR^ progeny arose from leaky somatic Cas9 activity in zygotes containing an intact *GSR* and the Cas9 (Fig. S1A). By further crossing GFP^-^ progeny to *Act-Cas9* and counting those that yielded GFP-positive F2, we derived the frequencies of intact *GSR*, NHEJ repair, and SSA repair in each case (Fig. S1A). These results revealed that SSA is the predominant repair pathway for *GSR* in the germline, in stark contrast to somatic tissues where NHEJ is as frequent as SSA (Fig. 1). This explains the high efficiency of the germline *GSR* assay even without a *Lig4* mutation.

**Figure 2.**
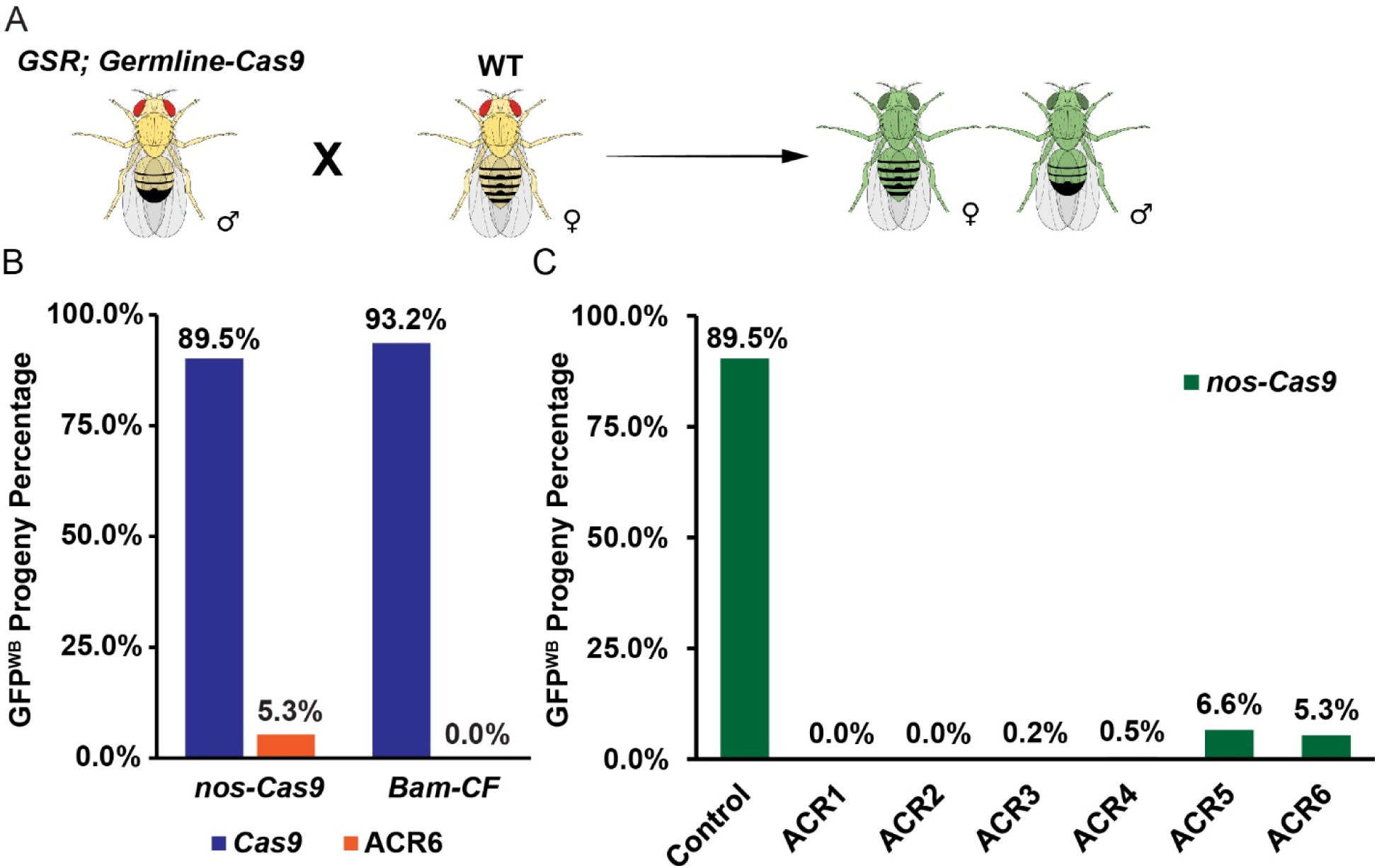
AcrIIA4-bearing balancers efficiently suppress germline Cas9 activity. (A) Scheme of germline GSR assay. (B) Frequencies of *GSR* activation by two different germline-Cas9 with and without the ACR6 *TM6B-AcrIIA4* balancer. n = total progeny number, *nos-Cas9* (n = 4607), *nos-cas9* with ACR6 (n = 3380), *Bam-CF* (n = 4693), *Bam-CF* with ACR6 (n = 1912). (C) Frequencies of *GSR* activation by *nos-Cas9* with different ACR balancers. n = total progeny number, ACR1 (n = 2179), ACR2 (n = 2060), ACR3 (n = 1920), ACR4 (n = 2600), ACR5 (n = 2270), ACR6 (n = 3380).

Next, we tested one *TM6B-AcrIIA4* balancer with both *nos-Cas9* and *bam-CF*. It robustly suppressed Cas9 activity, eliminating *bam-CF* activity and reducing *nos-Cas9*-mediated GSR activation to just 5.3% (Fig. 2B). Tests of all AcrII4 balancers (Table 1) against *nos-Cas9* showed that while all were effective, the *FM7c* (Chr. X) and *CyO* (Chr. II) balancers provided complete and near complete suppression, respectively, whereas the *TM6B* (Chr. III) balancers resulted in comparable levels of residual Cas9 activity (Fig. 2C). Thus, AcrIIA4 balancers can efficiently inhibit CRISPR/Cas9 in the *Drosophila* germline.

### AcrIIA4 balancers suppress somatic Cas9 activity and enable maternal-effect temporal control

A second application of AcrII4 balancers is to suppress somatic Cas9 activity in stocks, thereby preventing sickness or lethality caused by tissue-specific KO of essential genes. Cas9-mediated KO of essential genes, even in limited somatic tissues, can compromise animal health and stock maintenance. Acr balancers can suppress this activity, allowing the Cas9/gRNA combination to be maintained in stable and healthy stocks. To illustrate this, we combined a sensory neuron-specific *ppk-Cas9* with *gRNA-Nmnat*, targeting an enzyme required for NAD^+^ biosynthesis ^26^. This combination alone caused severe dendrite degeneration of class IV da (C4da) neurons as indicated by the dendritic marker *ppk-MApHS* ^27^ (Fig. 3A, 3B, 3E) due to activation of the conserved Wallerian degeneration pathway ^28^. However, when the *ppk-Cas9 gRNA-Nmnat* chromosome was balanced by *TM6B-AcrIIA4*, the animals showed wildtype (WT) dendrite morphology in larvae (Fig. 3C, 3E) and apparently normal fitness in adults. Thus, the *ppk-Cas9 gRNA-Nmnat* / *TM6B-AcrIIA4* strain can be maintained as a healthy, stable stock that, when crossed, conveniently introduces biallelic *Nmnat* mutations in the progeny’s sensory neurons.

**Figure 3.**
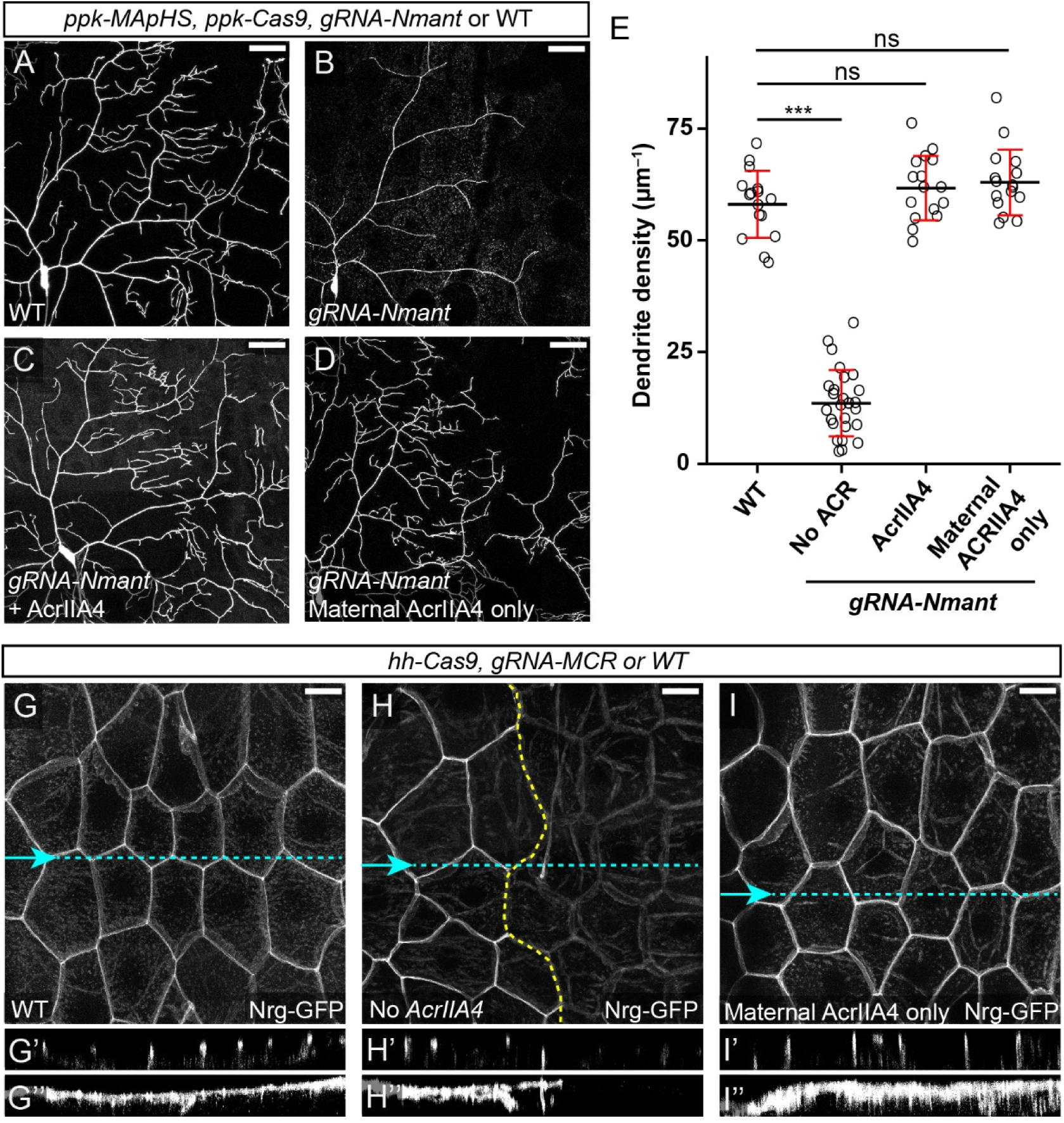
AcrIIA4 balancers suppress somatic Cas9 activity and enable maternal-effect temporal control. (A-D) C4da neurons expressing *ppk-Cas9* without gRNAs (A), with *gRNA-Nmnat* alone (B), with *gRNA-Nmnat* and the ACR6 balancer (C), with *gRNA-Nmnat* and maternal AcrIIA4 derived from ACR6 (D). C4da neurons were labeled by *ppk-MApHS.* (E) Dendrite density in indicated genotypes. n = number of neurons, WT (n = 16), *gRNA-Nmnat* only (n = 26), *gRNA-Nmnat* with AcrIIA4 (n = 16), *gRNA-Nmnat* with maternal AcrIIA4 only (n = 16). Black bar, mean; red bar, SD. One-way ANOVA and Tukey’s HSD test. ***p≤0.001, ns, not significance. (G-I”) The larval epidermis of *hh-Cas9*-expressing animals, visualized by the septate junction marker Nrg-GFP, without gRNAs (G-G”), with *gRNA-Mcr* (H-H”), and with *gRNA-Mcr* and AcrIIA4 derived from ACR6 (I-I”). Maximal projections of X/Y optical sections are shown in (G, H, and I). Single X/Z cross-sections along the blue dashed lines are shown in (G’, H’, and I’). Maximal projections of X/Z cross-sections are shown in (G”, H”, and I”). The border between the anterior and posterior hemisegments in (H) is indicated by the yellow dotted line. Scale bar, 50 µm for (A-D), 25 µm for (G-I).

Interestingly, when *ppk-Cas9 gRNA-Nmnat* / *TM6B-AcrIIA4* female parents were crossed to WT males to generate progeny lacking the Acr balancer, the *ppk-Cas9 gRNA-Nmnat* / *+* progeny did not exhibit any dendrite degeneration in late 3^rd^ instar larvae (Fig. 3D, 3E). This suggests that AcrIIA4 protein or mRNA, deposited into the oocyte from the *Tub-AcrIIA4* mother, is stable enough to inhibit Cas9 activity throughout embryonic and larval development. To further investigate this maternal-effect inhibition in a different cell type, we targeted Macroglobulin complement-related (Mcr), a septate junction protein essential for epithelium integrity ^29^. Combining *gRNA-Mcr* with *hh-Cas9* caused pre-pupal lethality and a loss of the septate junction marker Nrg-GFP ^30^ at apical junctions of epidermal cells in the posterior half of each segment (Fig. 3G-3H”), demonstrating efficient tissue-specific KO. In contrast, animals of the same genotype but derived from *TM6B-AcrIIA4* mothers showed normal epidermal junctions (Fig. 3I-3I”), confirming the potent, long-lasting inhibition by maternal AcrIIA4. Significantly, these animals still succumbed at the pre-pupal stage, indicating that the maternally supplied AcrIIA4 was eventually depleted or diluted, allowing Cas9 to become active at or after the larval-pupal transition.

Together, these results show that Acr balancers can be used not only to maintain stocks, but also as a powerful tool for temporal control, enabling studies of gene function during metamorphosis by suppressing Cas9 activity throughout all larval stages.

### Tissue-specific AcrIIA4 enables a “tissue-subtraction” strategy to refine Cas9 activity

A common problem with ts-CRISPR is leaky Cas9 activity in unintended tissues, which can confound phenotypic analysis ^9, 11, 14^. A possible solution is to express Acr specifically in the unintended tissue to “subtract” Cas9 activity there. To build tools for this “tissue-subtraction” strategy, we established two methods for generating tissue-specific AcrII4 lines: (1) When a tissue-specific enhancer is available, it can be introduced into an AcrIIA4 expression vector through Gateway LR cloning (Fig. S2A). We generated an epidermal specific *shot-AcrIIA4* and a glial specific *repo-AcrIIA4* this way. (2) When a simple enhancer is not sufficient or available, we use genomic regions (e.g., 5’ and 3’ flanking sequences) of tissue-specific genes to drive AcrIIA4 expression (Fig. S2B). A neuronal-specific *RabX4-AcrIIA4* and a germline-specific *nos-AcrIIA4* were made this way.

To test the tissue-subtraction strategy, we chose *sop-Cas9*, which is expressed in the progenitor cells of peripheral sensory neurons but also shows significant leaky activity in surrounding epidermal cells ^8^. We hypothesized that adding the epidermal *shot-AcrIIA4* would remove the leaky Cas9 activity, resulting in a cleaner, neuron-specific Cas9 (Fig. 4A). By itself, *sop-Cas9* activated GSR in a 10.9% area of the larval body wall, including both sensory neurons (yellow arrowheads) and many epidermal cells (green arrowheads) (Fig. 4B, 4D). The addition of *shot-AcrIIA4* abolished nearly all GFP in epidermal cells without apparent effects on neuronal labeling, reducing total GSR activation to a 3.4% area (Fig. 4C, 4D). Thus, tissue-specific AcrIIA4 can be used to “clean up” leaky Cas9 lines and improve the precision of ts-CRISPR.

**Figure 4.**
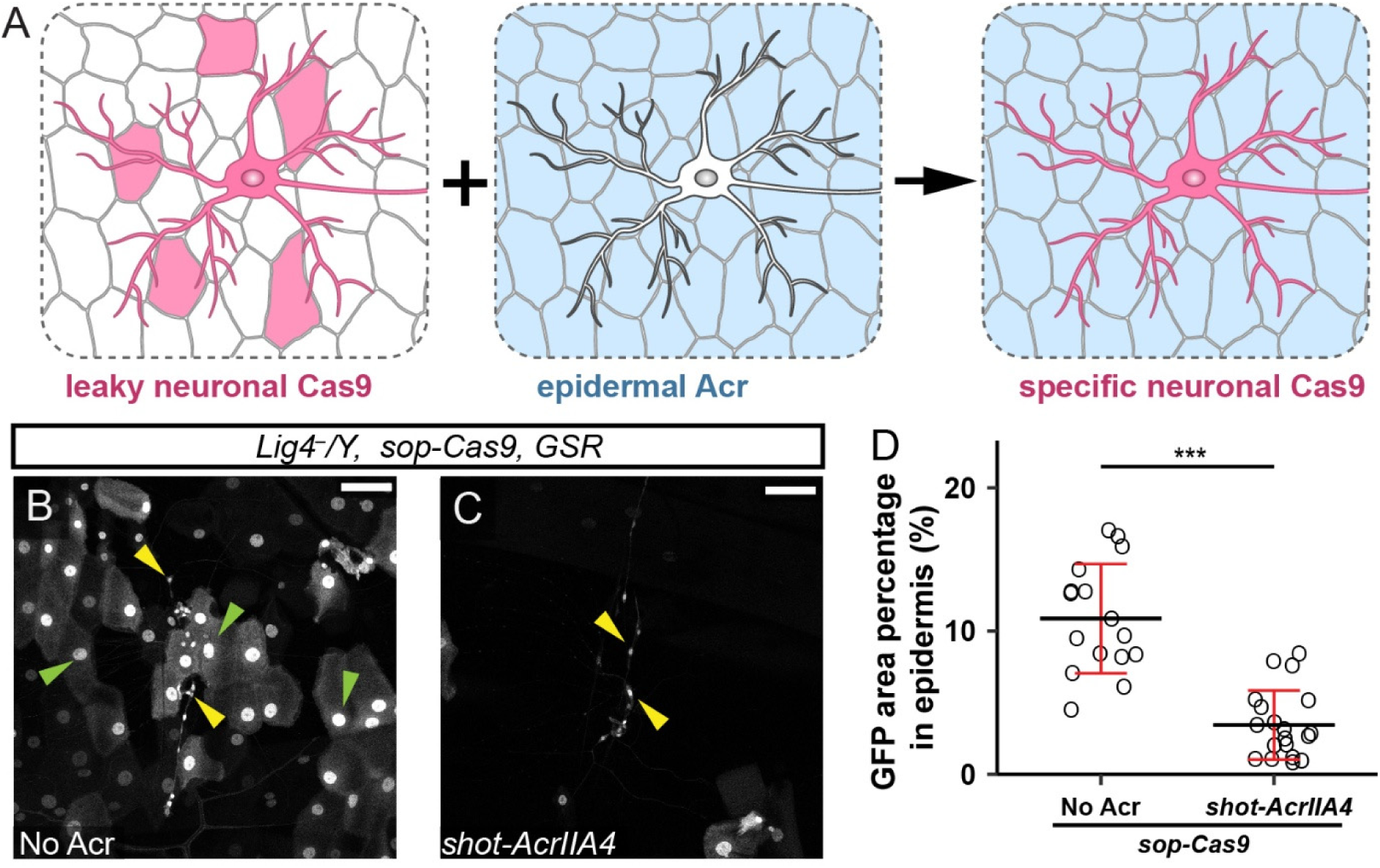
Tissue-specific AcrIIA4 enables a “tissue-subtraction” strategy to refine Cas9 activity. (A) Scheme of how an epidermis-specific Acr refines the activity of a leaky neuronal Cas9. (B and C) Activity pattern of *sop-Cas9* visualized by *GSR* activation in the *Lig4^-^* hemizygote background without Acr (B) and with *shot-AcrIIA4* (C). Peripheral neurons are indicated by yellow arrowheads, and epidermal cells are indicated by green arrowheads. Scale bar, 50 µm. (D) Percentage of GFP-positive area in the larval epidermis. n = number of segments, *sop-Cas9* (n = 16), *sop-Cas9* with *shot-AcrIIA4* (n = 19). Black bar, mean; red bar, SD. Student’s t-test. ***p≤0.001.

### AcrIIA4 requires specific temporal windows to prevent Cas9 activity

Because Cas9-induced mutations are irreversible, we next sought to determine the temporal window in which AcrIIA4 must be expressed to effectively suppress Cas9. First, we tested simultaneous expressions of Cas9 and AcrIIA4 driven by identical promoters. *repo-Cas9* on its own caused widespread (81.9%) GSR activation in glial cells of the larval brain (Fig. 5A, 5C). The addition of *repo-AcrIIA4* reduced GSR activation to 4.5%, leaving only rare GFP-positive glial cells (Fig. 5B, 5C). Similarly, *shot-Cas9* activated GSR in an 81% epidermal area, but activity was reduced to merely 8.5% after adding *shot-AcrIIA4* (Fig. 5D-5F). These data suggest that, when expressed at the same time, AcrIIA4 can dominantly suppress Cas9 activity.

**Figure 5.**
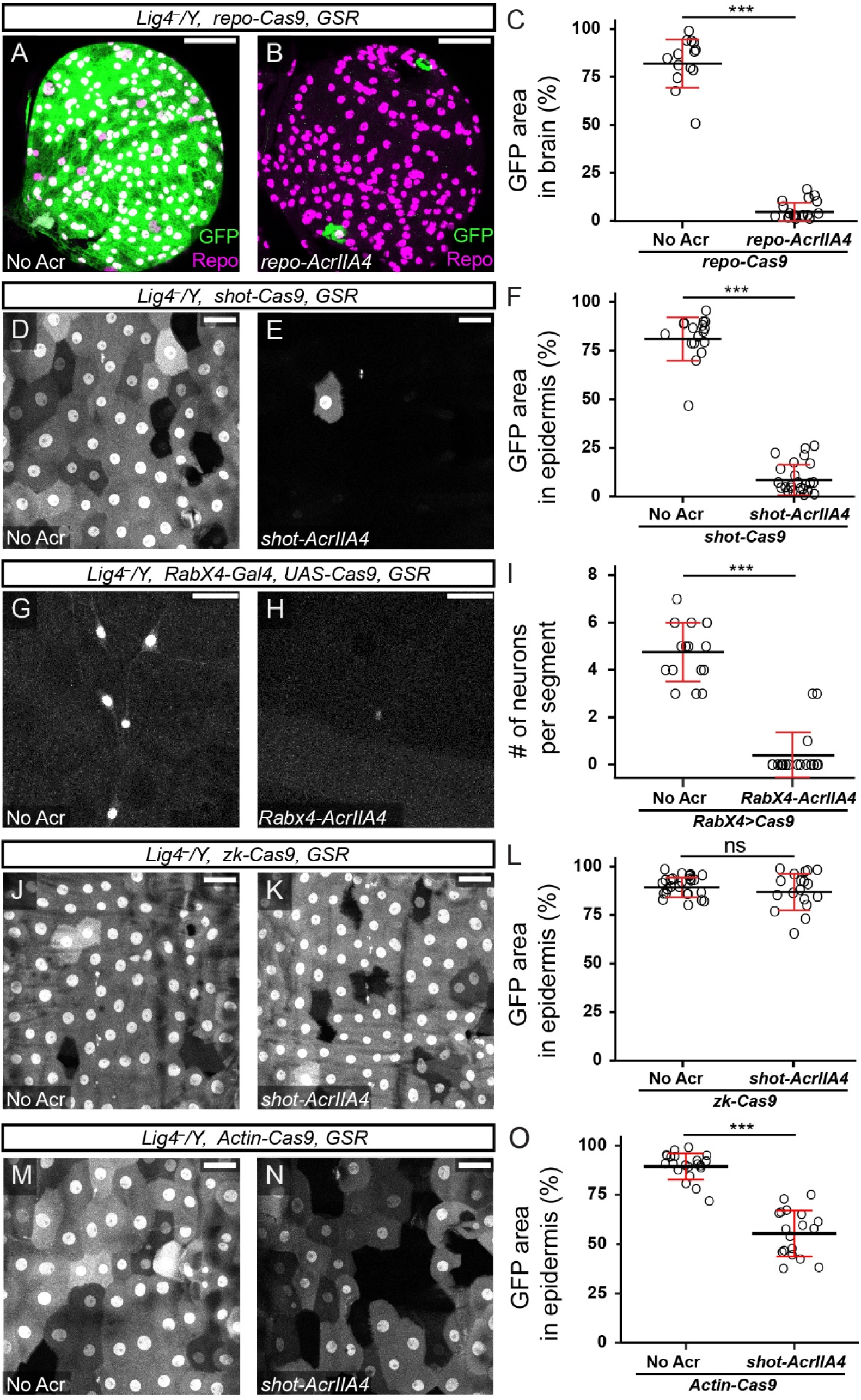
AcrIIA4 requires specific temporal windows to prevent Cas9 activity. (A and B) Activity pattern of *repo-Cas9* without Acr (A) and with *repo-AcrIIA4* (B). Glial nuclei were labeled by Repo staining. (C) GFP-positive area in the larval brain. n = number of brain lobes, *repo-Cas9* (n = 14), *repo-Cas9* with *repo-AcrIIA4* (n = 18). (D and E) Activity pattern of *shot-Cas9* without Acr (D) and with *shot-AcrIIA4* (E). (F) GFP-positive area in the larval epidermis. n = number of segments, *shot-Cas9* (n = 17), *shot-Cas9* with *shot-AcrIIA4* (n = 24). (G and H) Activity pattern of *RabX4-Gal4 UAS-Cas9* without Acr (G) and with *RabX4-AcrIIA4* (H). (I) The number of labeled peripheral neurons in each segment. n = number of segments, *RabX4-Gal4 UAS-Cas9* (n = 16), *RabX4-Gal4 UAS-Cas9* with *RabX4-AcrIIA4* (n = 18). (J and K) Activity pattern of *zk-Cas9* without Acr (J) and with *shot-AcrIIA4* (K). (L) GFP-positive area in the larval epidermis. n = number of segments, *zk-Cas9* (n = 26), *zk-Cas9* with *shot-AcrIIA4* (n = 18). (M and N) Activity pattern of *Actin-Cas9* without Acr (M) and with *shot-AcrIIA4* (N). (O) GFP-positive area in the larval epidermis. n = number of segments, *Actin-Cas9* (n = 21), *Actin-Cas9* with *shot-AcrIIA4* (n = 11). In all plots, black bar, mean; red bar, SD. Student’s t-test. ***p≤0.001, ns, not significance. Scale bar, 50 µm for (A, B, D, E, J, K, M and O), 25 µm for (G and H). In all experiments, Cas9 activity was visualized by *GSR* activation in the *Lig4^-^*hemizygous mutant background.

We next asked if AcrIIA4 expressed early could suppress Cas9 expressed later, particularly if Cas9 was expressed at a much higher level. For this purpose, we used the UAS/Gal4 binary system (*RabX4-Gal4*>*UAS-Cas9*) to express a relatively high level of Cas9 in neurons and the enhancer-fusion *RabX4-AcrIIA4* transgene to express a relatively low level of AcrIIA4. *RabX4-Gal4 > UAS-Cas9* is expected to express Cas9 later than the direct enhancer-driven *RabX4-AcrIIA4* due to the delay from Gal4 transcription and translation. While *RabX4-Gal4*>*UAS-Cas9* alone activated GSR in 3-7 larval sensory neurons in every segment (Fig. 5G, 5H), addition of *RabX4-AcrIIA4* abolished GSR activation in 83% of segments and led to fewer GFP neurons in the remaining outliers (Fig. 5H, 5I). This demonstrates that AcrIIA4, even when expressed early and at a likely lower level, can potently suppress later, high-level Cas9 activity.

Lastly, we asked if a late AcrII4 transgene can suppress Cas9 expressed early. For this purpose, we first tested *shot-AcrIIA4* with *zk-Cas9*. The *zen-Kruppel* enhancer in *zk-Cas9* becomes active in the dorsal ectoderm of the late blastoderm embryo ^31, 32, 33^, long before the *shot*/R38F11 enhancer in *shot-AcrIIA4* becomes active in the larval epidermis ^34^. As expected, given the irreversible nature of Cas9-induced mutations, *shot-AcrIIA4* had no effect; both *zk-Cas9* alone and in combination with *shot-AcrIIA4* showed near-complete GSR activation in the epidermis (Fig. 5J-5L). Next, we tested *shot-AcrIIA4* (the late Acr) with *Actin-Cas9* (an early and continuous Cas9). Interestingly, adding *shot-AcrIIA4* reduced epidermal GSR activation by *Actin-Cas9* from 89% to 55% (Fig. 5M-5O). We posit that *Actin-Cas9* is continuously active from the blastoderm through larval stages; the late-expressing *shot-AcrIIA4* cannot reverse mutations that already occurred but can stop *Actin-Cas9* from inducing new GSR activation during the larval stage, resulting in partial suppression.

Together, the above data show that AcrIIA4 can efficiently suppress Cas9 activity when expressed earlier than or at the same time as Cas9, but it can only partially suppress a continuous Cas9 if expressed later, highlighting the importance of expression timing.

### Germline- and soma-specific AcrIIA4 tools offer flexible control of germline Cas9 reagents

A major challenge in ts-CRISPR is the bidirectional leakiness between somatic and germline compartments: many somatic Cas9 drivers are leaky in the germline, and germline drivers are leaky in the soma. To solve this, we developed AcrIIA4 tools to segregate Cas9 activity between these two compartments.

We first tested *nos-AcrIIA4*, our germline-specific Acr tool. It was tested with the germline-specific *nos-Cas9*, a ubiquitous *Tub-Cas9* ^35^, and *hh-Cas9*, a somatic epithelial driver ^8^. As expected, *nos-Cas9* and *Tub-Cas9* parents produced => 90% GFP^WB^ progeny (Fig. 6A, 6B). Surprisingly, the somatic *hh-Cas9* driver also produced > 1/3 GFP^WB^ progeny (Fig. 6A, 6B), revealing significant germline leakiness. Importantly, adding *nos-AcrIIA4* to the parents suppressed germline activity from all three drivers, reducing GFP^WB^ progeny to <3% for *nos-Cas9* and *Tub-Cas9*, and eliminating it for *hh-Cas9* (Fig. 6A, 6B). Thus, *nos-AcrIIA4* is a powerful tool for preventing germline transmission of mutations from somatic or leaky Cas9 drivers.

**Figure 6.**
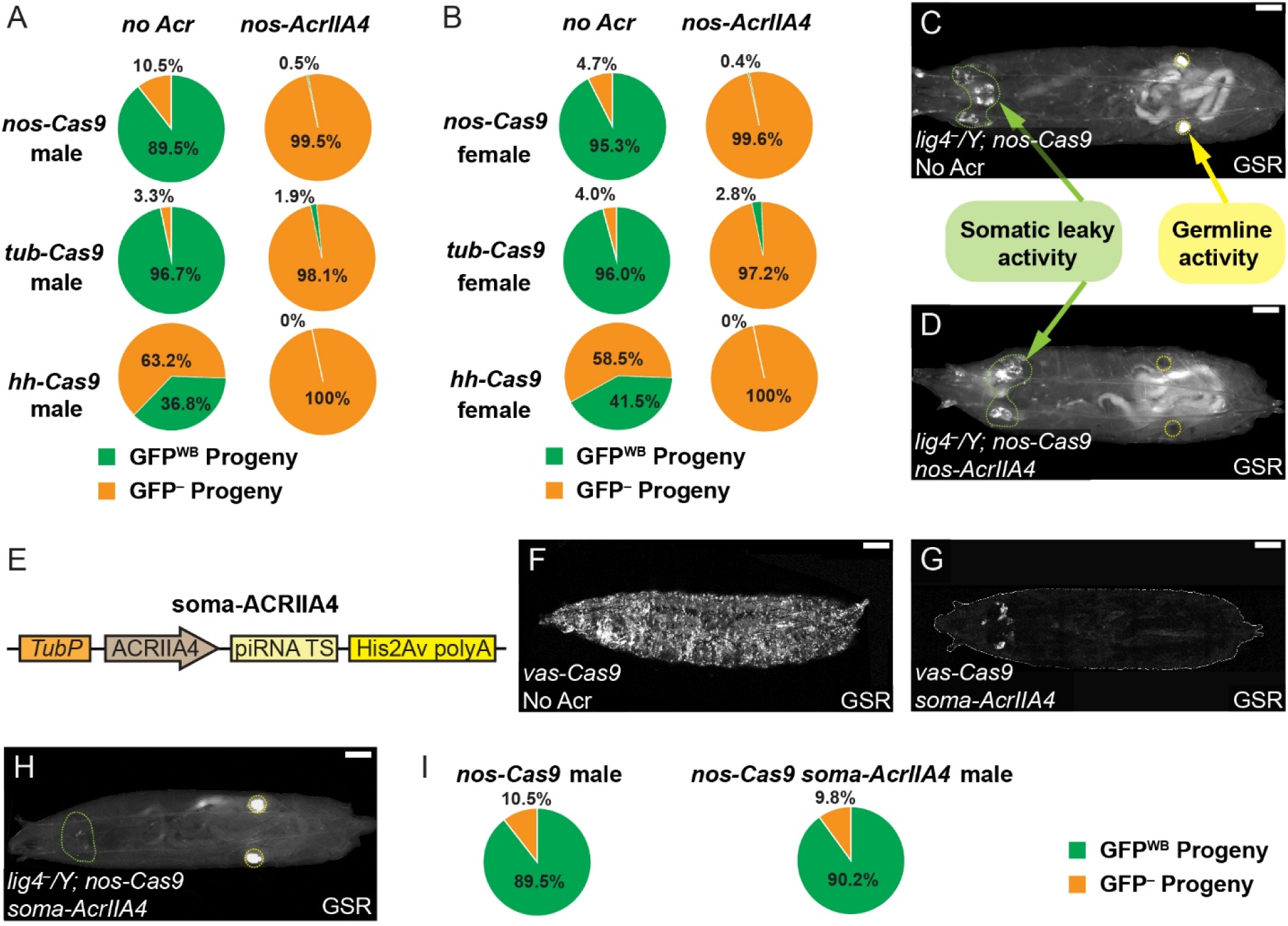
Germline- and soma-specific AcrIIA4 tools offer flexible control of germline Cas9 reagents. (A and B) Pie charts of germline activity of *nos-Cas9, tub-Cas9*, and *hh-Cas9* in males (A) and females (B) with and without *nos-AcrIIA4* using the germline GSR assay. n = total progeny number, *nos-Cas9* male (n = 4607), *nos-cas9* male with *nos-AcrIIA4* (n = 1480), *tub-Cas9* male (n = 1220), *tub-cas9* male with *nos-AcrIIA4* (n = 1160), *hh-Cas9* male (n = 950), *hh-cas9* male with *nos-AcrIIA4* (n = 920), *nos-Cas9* female (n = 5690), *nos-cas9* female with *nos-AcrIIA4* (n = 1440), *tub-Cas9* female (n = 990), *tub-cas9* female with *nos-AcrIIA4* (n = 1010), *hh-Cas9* female (n = 265), *hh-cas9* female with *nos-AcrIIA4* (n = 280). (C and D) Activity pattern of *nos-Cas9* in the whole larval body without Acr (C) and with *nos-AcrIIA4* (D). Green arrows and green dashed lines indicate the leaky somatic activity of *nos-Cas9* in the head region; the yellow arrow and yellow dashed lines indicate the locations of gonads. (E) The construct design of *soma-AcrIIA4*. TS, target sequence. (F and G) Activity pattern of *vas-Cas9* in the whole larval body without Acr (F) and with *soma-AcrIIA4* (G). (H) Activity pattern of *nos-Cas9* in the whole larval body with *soma-AcrIIA4*. The green dashed line indicates the head region where the leaky somatic activity of *nos-Cas9* is expected; yellow dashed lines indicate the locations of gonads. (I) Pie charts showing germline activity of *nos-Cas9* with or without *soma-AcrIIA4.* The dataset of *nos-Cas9* without *soma-AcrIIA4* is the same as in (A). n = total progeny number, *nos-Cas9* male (n = 4607), *nos-cas9* male with *nos-AcrIIA4* (n = 494). For all images, scale bar, 500 µm. The activity of *nos-Cas9* was visualized by GSR activation in the *Lig4^-^* hemizygous mutant background. The activity of *vas-Cas9* was visualized by GSR activation in the WT *Lig4* background.

Conversely, when examining *nos-Cas9* (a germline driver) in larvae, we observed the expected GSR activation in the gonad, but also significant somatic leakiness, particularly in the head region (Fig. 6C). As expected, *nos-AcrIIA4* eliminated the germline labeling but not the somatic leaky labeling (Fig. 6D), confirming that *nos-AcrIIA4* itself is germline-specific. To create the reciprocal tool, a soma-specific Acr (*soma-AcrIIA4*), we added piRNA targeting sequences to the 3’ untranslated region (UTR) of the ubiquitous *Tub-AcrIIA4* construct (Fig. 6E). Because piRNAs mediate RNAi only in the germline ^36^, this should restrict AcrIIA4 expression to somatic tissues. As predicted, *soma-AcrIIA4* abolished most somatic GSR activation from the ubiquitous *vas-Cas9* driver ^6, 37^ (Fig. 6F, 6G). Crucially, *soma-AcrIIA4* provided the desired reciprocal result to *nos-AcrIIA4*: it eliminated the somatic leaky activity of *nos-Cas9* while having no effect on its intended gonadal activity (Fig. 6H) or its ability to generate GFP^WB^ progeny (Fig. 6I).

Together, *nos-AcrIIA4* and *soma-AcrIIA4* constitute a versatile toolkit for separating germline and somatic activities of Cas9 lines, significantly improving the precision of *Drosophila* CRISPR applications.

## DISCUSSION

In this study, we explored the potential of Acrs for enhancing the precision and versatility of *in vivo* CRISPR/Cas9 applications in *Drosophila*. We first identified AcrIIA4 as the most potent AcrIIA protein for suppressing Cas9 activity in *Drosophila* somatic tissues and then determined the temporal rules for its efficient application. Based on these findings, we developed three classes of Acr tools to solve common challenges in ts-CRISPR experiments: (1) Acr-carrying balancers for stably maintaining strains containing both Cas9 and gRNA transgenes and, via maternal effect, for delayed knockouts; (2) tissue-specific AcrIIA4 lines for eliminating leaky Cas9 activity in non-target tissues using a “tissue-subtraction” strategy; (3) germline- and soma-specific AcrIIA4 lines for segregating Cas9 activities between these two compartments. Lastly, we offered two approaches to generate new tissue-specific AcrIIA4 transgenes. These tools are important additions to the CRISPR arsenal in *Drosophila* and will enable more sophisticated genetic studies and functional analysis of genes.

### *In vivo* properties of AcrIIA4

AcrIIA4, AcrIIA5, and AcrIIA16 are all known potent inhibitors of SpyCas9, the predominant nuclease used for *Drosophila* genome editing, though they operate via different mechanisms. AcrIIA4 binds to the PAM-interacting domain of sgRNA-loaded Cas9, preventing PAM recognition and locking it in an inactive conformation ^17, 21^. AcrIIA5 contains an intrinsically disordered region that interacts with Cas9 surfaces involved in DNA cleavage, impairing catalysis without blocking DNA binding ^22^. In contrast, AcrIIA16 interacts with Cas9 and degrades the gRNA to interfere with the assembly of the Cas9–sgRNA complex ^23, 38^. While previous studies in transfected human cells suggested that the three proteins have comparable inhibitory potencies ^38^, their relative efficacy in a whole-animal context was unknown. In this study, we show that ubiquitously expressed AcrIIA4 is markedly more potent than AcrIIA5 and AcrIIA16 in *Drosophila* and was the only one capable of fully suppressing *hh-Cas9* activity. This finding demonstrates the necessity of *in vivo* comparison before choosing an Acr for animal applications, as cell culture potency does not necessarily translate to whole-organism efficacy.

Our further analyses revealed important *in vivo* properties of AcrIIA4. Temporally, to completely block Cas9 activity in a given cell lineage, AcrIIA4 must be expressed at the same time as or earlier than Cas9. This rule is explained by the irreversible nature of Cas9-induced mutations: once a gene is mutated, a later-expressing AcrIIA4 cannot restore it. However, our finding that late-expressing AcrIIA4 partially suppresses a continuously-expressed Cas9 (like *Actin-Cas9*) suggests that AcrIIA4 can stop new Cas9 activity, even if it cannot reverse mutations that have already occurred.

Furthermore, our results revealed an *in vivo* inhibitory capacity not predicted by *in vitro* studies. While *in vitro* studies demonstrated stoichiometric inhibition of Cas9 by AcrIIA4 ^17, 21^, our results instead show that AcrIIA4 can functionally suppress an apparent excess of Cas9 *in vivo*. First, an enhancer-driven AcrIIA4 (*RabX4-AcrIIA4*) nearly completely suppressed Gal4/UAS-driven Cas9, which is expected to produce much higher protein levels in the same tissue. Second, maternally inherited AcrIIA4 suppressed highly expressed Cas9 until the end of the larval stage. These results indicate that AcrIIA4 a highly stable protein *in vivo*, allowing a lower-level or earlier-expressed Acr to effectively neutralize a subsequent, larger pool of Cas9 protein over long developmental periods.

### Novel CRISPR applications enabled by Acr

Beyond defining its *in vivo* properties, we demonstrated how AcrIIA4 provides a powerful second layer of control to make ts-CRISPR more precise, flexible, and robust.

First, AcrIIA4 can be used to improve the precision of ts-CRISPR. Because Cas9 permanently modifies DNA, even transient leaky expression in non-target tissues can cause confounding phenotypes. Our tissue-specific AcrIIA4 tools directly solve this with a “subtraction” strategy. For example, *shot-AcrIIA4* removes leaky Cas9 activity from epidermal cells and significantly improves the tissue-specificity of the neuronal precursor *SOP-Cas9*. Similarly, our germline- and soma-specific AcrIIA4 tools faithfully segregate Cas9 activity between these two compartments. *nos-AcrIIA4* can preserve somatic KO phenotypes while preventing germline mutations that would otherwise destabilize the stock. Conversely, *soma-AcrIIA4* removes confounding somatic leakiness from germline drivers like *nos-Cas9*, making germline KO experiments cleaner and more viable. Importantly, *soma-AcrIIA4* significantly improves the Gal4-to-Cas9 conversion workflow ^11^. This workflow uses a germline Cas9 to convert a Gal4 line, but leaky somatic activity from the germline Cas9 often obscures the resulting “converted” pattern. By recombining *soma-AcrIIA4* with the germline Cas9, this background activity is eliminated, making screening for convertants far more efficient.

Second, Acr balancers enable much more flexible ts-CRISPR. By introducing an Acr balancer to a strain carrying both a tissue-specific Cas9 and gRNAs, the *Tub-AcrIIA4* on the balancer serves two purposes: it prevents germline mutations that would inactivate the gRNA, and it suppresses somatic activity that could cause sickness or lethality. These two features make long-term maintenance of the stock possible. With such strains, one can introduce Gal4-independent, biallelic, and tissue-specific KO of a gene of interest into any genetic background by a simple cross. Such strains can be used to greatly expedite gene discovery, for example, in genome-wide modifier screens.

Third, the maternal effect of Acr balancers provides a novel method for temporal control of Cas9 activity. In progeny from a *Tub-AcrIIA4* (balancer) mother, Cas9 activity is blocked throughout embryonic and larval development. This creates a temporal window to study the roles of essential genes during metamorphosis and adulthood, bypassing pleiotropic effects or lethality from earlier knockouts.

### Future improvements of Acr tools

This study establishes a foundation for *in vivo* Acr application. Besides the approaches described above, Acr proteins may be engineered for even more sophisticated control. For example, the long-lasting stability of AcrIIA4, while useful for suppression of zygotic Cas9 by maternal AcrIIA, could interfere with experiments requiring sequential CRISPR activity. Destabilized AcrIIA4 variants (e.g. fused to a PEST domain) may overcome this limitation. Second, a temperature-sensitive AcrIIA4 variant would allow for reversible temporal control. Third, drug-dependent AcrIIA4 transgenes could make ts-CRISPR inducible rather than suppressive. Lastly, recent optogenetic AcrIIA4 tools ^39^ could be adapted for *Drosophila* to provide precise spatiotemporal control of CRISPR activity with light.

In summary, our work establishes Acr as a powerful extension to existing *Drosophila* genetic tools, resolving major challenges in ts-CRISPR and opening new avenues for precise, flexible, and interpretable genetic studies. The benefits of Acr demonstrated here—spatial and temporal refinement of CRISPR, stable coexistence of Cas9 and gRNAs, and novel experimental designs—may be applicable to other CRISPR systems, such as CRISPR activation (CRISPRa), CRISPR interference (CRISPRi), and base editors. The *in vivo* principles of AcrIIA4 learned in this study may also inform the deployment of Acr technology in other animal models.

## MATERIALS AND METHODS

### Fly Stocks and Husbandry

See the Key Resource Table (S1 Table) for details of fly stocks used in this study. Most fly lines were either generated in the Han lab or obtained from the Bloomington *Drosophila* Stock Center. All flies were grown on standard yeast-glucose medium, in a 12:12 light/dark cycle, at 25°C unless otherwise noted. Virgin males and females for mating experiments were aged for 3-5 days.

To acquire hemizygotes of *Lig4* null mutant flies, Cas9 flies with or without AcrIIA4 were crossed to *Lig4^-/-^; GSR* flies. Male progenies were then selected for living image. *vas-Cas9*, with or without *soma-AcrIIA4*, was tested with *GSR* in the WT *Lig4* background. To acquire maternal AcrIIA4 only animals, *ppk-Cas9 gRNA-Nmant* or *hh-Cas9 gRNA-Mcr* flies were balanced with *TM6b-tub-AcrIIA4.w+*. The virgins were then crossed with wild type males; in the progeny, non-*Tb* larvae were selected for live imaging.

### Molecular Cloning

#### Tub-AcrIIA constructs

A DNA fragment containing the coding sequence (CDS) of AcrIIA4 followed by the nuclear localization signal (NLS) of SV40 was synthesized by Integrated DNA Technologies, Inc. (IDT). The *αTub84B* promoter was PCR-amplified from pENTR221-tub ^35^ using oligos ctcggcaacagcatgctgcagAGATCTTGCACAGGTCCTGTTCGATAAC and GGATCCctgtggatgaggaggaagggaaaac. Both fragments were assembled into the pAPIC vector ^40^ through NEBuilder HiFi DNA assembly (New England Biolabs, Inc) to make pAPIC-tub-AcrIIA4. The pAPIC vector was obtained by digesting pAPIC-ppk-CD4-tdGFP(HCH) ^40^ with PstI and PacI. Next, the AcrIIA4 CDS in pAPIC-tub-AcrIIA4 was replaced by those of AcrIIA5 and AcrIIA16 through BamHI and NheI digestions and assembly reactions with DNA fragments synthesized by IDT. This resulted in the expression constructs pAPIC-tub-AcrIIA5 and pAPIC-tub-AcrIIA16. To make pAPIC-tub-AcrIIA4(RFP), a 3xP3-DsRed cassette was PCR-amplified from pXL-BACII-LoxP-3xPDsRed-LoxP (Addgene 26852) using oligos cattttttttttttactgcactggatCCCACAATGGTTAATTCGAGC and gctggcgcaggatattagatagtttggacaaaccacaactagaatg and assembled into EcoRV-digested pAPIC-tub-AcrIIA4.

#### AcrIIA4 destination vector

The AcrIIA4 CDS followed by SV40 NLS was PCR-amplified using pAPIC-tub-AcrIIA4 as the template and was used to replace the Cas9 CDS in pDEST-APIC-Cas9 ^8^ between XhoI and PacI sites. This resulted in pDEST-APIC-AcrIIA4.

#### AcrIIA4 expression vector

*shot-AcrIIA4 and repo-AcrIIA4* expression vectors were generated by Gateway LR reactions (Thermo Fisher Scientific) using entry vectors pENTR221-R38F11 ^28^ and pENTR-repo ^11^ and the destination vector pDEST-APIC2-AcrIIA4.

#### RabX4-AcrIIA4 and nos-AcrIIA4

pAPIC-Rabx4-AcrIIA4 and pAPIC-nos-AcrIIA4 were generated by replacing the *αTub84B* promoter and the *His2Av* 3’UTR and polyA in pAPIC-tub-AcrIIA4 via two sequential steps of NEBuilder HiFi DNA assembly cloning. The 5’ and 3’ sequences flanking the coding exons of *RabX4* and *nos* were PCR-amplified from the genomic DNA of *w^1118^*. The oligos used were atcaattgtgctcggcaacagcatgctgcgcatgggatactatgaaac and GATCATTGATGTTGCCCATGGATCCATTCATgttgacggagctggcggagt for *RabX4* 5’; CAAAGAAAAAGCGAAAGGTCTAAtctagatactcagcgatggattacgatg and aacgcacacttattacgtgACTAGTgaggacgaccatatttgcagggccac for *RabX4* 3’; ttgtgctcggcaacagcatgcattgaaagcttcgaccgttttaacc and GATCATTGATGTTGCCCATGGATCCATTCATggcgaaaatccgggtcgaaag for *nos* 5’; and CAAAGAAAAAGCGAAAGGTCTAAtctagagagggcgaatccagctctg and aacgcacacttattacgtgACTAGTACAGTGATCTTACCGATGGCATCTTC for *nos* 3’.

#### Soma-AcrIIA4

A 1312-bp DNA fragment containing piRNA targeting sequences was synthesized by IDT and assembled into KpnI digested pAPIC-tub-AcrIIA4 to make pAPIC-soma-AcrIIA4. The fragment was made of a 252 bp consensus sequence at Su(Ste) cluster ^41^ and a 1000-bp sequence from piRNA cluster 20A (personal communication with Julius Brennecke and Ralf Jansen).

#### gRNA-Mcr

Three gRNA targeting sequences (AGGAGGCTATCAGGACAACG, GTATCAGGACAGCAGCTACG, and ACCTCGAAGAAGGATCCCTGTGG) were cloned into pAC-U61-SapI-Rev using protocols previously described ^8^. The three gRNA targeting sequences are under control of three *U6* promotors (U6:2, U6:2, and U6:3, respectively).

Injections were carried out by Rainbow Transgenic Flies (Camarillo, CA 93012 USA) or Genetivision (Stafford, TX 77477) to transform flies through φC31 integrase-mediated integration into attP docker sites.

### Tub-AcrIIA4 mobilization

*Tub-AcrIIA4* was mobilized and inserted onto *TM6B* using the transposase chromosome *CyO Delta2-3*. *Tub-AcrIIA4(RFP)* was mobilized and inserted onto *FM7c*, *CyO*, and *TM6B* using *TMS Delta2-3*. *Tub-AcrIIA4* or *Tub-AcrIIA4(RFP)* was combined with the *Delta2-3* chromosome and the target balancer in the same flies through sequential crosses. The flies were then crossed to *w^1118^* to remove the *Delta2-3* chromosome and the original *AcrIIA4* transgene. The remaining target chromosomes were then screened based on eye color (for *Tub-AcrIIA4*) or fluorescence (for *Tub-AcrIIA4(RFP)*).

### Germline *GSR* assay

Flies containing *GSR* and the test Cas9, with or without the test AcrIIA4, were crossed to wildtype flies. The progeny were scored as 3^rd^ instar larvae for the numbers of total larvae, whole-body GFP-positive (GFP^WB^) larvae, sporadic-and-random GFP (GFP^SR^) larvae, and GFP-negative (GFP^-^) larvae. The frequency of SSA repair was calculated as # GFP^WB^ ÷ # total larvae × 2 × 100%. 2 was multiplied due to heterozygosity of *GSR* in the test parents.

For Figure S1, *GSR* in the test parents was balanced by *CyO Tb* so that we could distinguish their *GSR*-carrying progeny (i.e. non-*Tb*). To calculate the frequency of intact *GSR* in F1, GFP^-^ non-*Tb* progeny were recovered as larvae and later crossed individually to *Actin-Cas9*. The number of F1 crosses that produced GFP-positive (GFP^+^) F2 larvae were counted as intact *GSR*. Thus, the frequency of intact *GSR* was calculated as (# GFP^SR^F1 + # GFP^+^) ÷ # non-*Tb*F1 × 100%, and the frequency NHEJ was calculated as # GFP^-^ ÷ # non-*Tb*F1 × 100%.

### Live Imaging

Live imaging of larval epidermal cells, sensory neurons was performed as previously described ^42^. Animals were collected at 96 (for late third larvae) or 120 hours AEL (for wandering third instar larvae) hours after egg-laying (AEL) and mounted in glycerol on a slide with vacuum grease as a spacer. Animals were imaged using a Leica SP8 confocal microscope with a 40X NA1.3 oil objective, pinhole size 2 airy units, and a z-step size of 1 µm. For epidermis, images were taken at the dorsal midline of A2 and A3 segments. For C4da neurons, images were taken from A2 to A3 hemi-segments. The live imaging of the whole larvae was done either using a Leica SP8 confocal microscope with a 10X air objective, pinhole size 3.5 airy units, and a Z step of 7 µm or a Nikon SMZ18 stereo microscope. Confocal images were taken using the tile function, which automatically stitched the images into a whole-large image. Nrg-GFP localization in the larval epidermis (Figure 3G-I) was imaged using a Leica SP8 confocal microscope with a 63X 1.4 oil objective, pinhole size 1 airy units, and a z-step size of 0.2 µm.

### Immunohistochemistry

Larval dissections were performed as described previously ^8^. Briefly, wandering third instar larvae were dissected in a small petri dish filled with cold phosphate-buffered saline (PBS). The anterior half of the larva was inverted. Trachea and gut were removed. Larval brains were then transferred to 4% formaldehyde in PBS and fixed for 20 minutes at room temperature. Larval brains were rinsed and washed at room temperature in PBS with 0.2% Triton-X100 (PBST) after fixation. The samples were then blocked in PBST with 5% normal donkey serum (NDS) for 1 hour before incubating with mouse anti-Repo antibody 8D12 (1:50 dilution) in the blocking solution. Brains were stained for 2 hours at room temperature. After additional rinsing and washing, the samples were incubated with donkey anti-mouse antibody conjugated with Cy5 (1:400) for 2 hours at room temperature. The samples were then rinsed and washed again before mounting and imaging.

### Image Analysis and Quantification

Image analyses were conducted in Fiji/ImageJ. To compare the ratio of GFP-labeled epidermis, cells were detected based on thresholds to generate masks. The ratio was then measured as the area of the mask divided by the area of the region of interest.

The tracing and measurement of da neuron dendrites were done as previously described ^42^. Briefly, dendrites were segmented using local thresholding. The segments were then converted into single-pixel-width skeletons. The total length of skeletons was calculated based on pixel distance. Normalized dendrite density was calculated as dendritic length (μm)/area of ROI (μm^2^).

### Statistical Analysis

We first confirmed that the dependent variables were normally distributed using Shapiro-Wilk tests and that there was approximately equal variance across groups using Levene’s tests. One-way analysis of variance (ANOVA) with Tukey’s HSD test was then used for statistical comparison among groups. For additional information on the number of samples, see figure legends. R studio was used for all statistical analyses and all plots.

## ACKNOWLEDGMENTS

We thank Developmental Studies Hybridoma Bank (DSHB) for antibodies; Tzumin Lee and Bloomington *Drosophila* Stock Center for fly stocks; Addgene for plasmids; Julius Brennecke, Ralf Jansen, Mariana Wolfner, Norbert Perrimon, and Tzumin Lee for advice; Rhiannon Clements and Elizabeth Korn for helping to establish and test transgenes; Mariana Wolfner for feedback on the manuscript. This work was supported by an NIH grant (R24OD031953) awarded to C.H..

## AUTHOR CONTRIBUTIONS

**Conceptualization:** Chun Han, Yifan Shen

**Data curation:** Yifan Shen, Ann Yeung

**Formal analysis:** Yifan Shen, Ann Yeung

**Funding acquisition:** Chun Han

**Investigation:** Yifan Shen, Michael Sheen, Ann Yeung, Xinchen Chen

**Methodology:** Chun Han, Yifan Shen

**Resources:** Bei Wang, Zixian Huang, Claire Ho, Zachary Lakkis

**Supervision:** Chun Han, Yifan Shen

**Validation:** Yifan Shen

**Visualization:** Yifan Shen

**Writing – original draft:** Chun Han, Yifan Shen

**Writing – review & editing:** Chun Han, Yifan Shen

---

**Figure S1.**
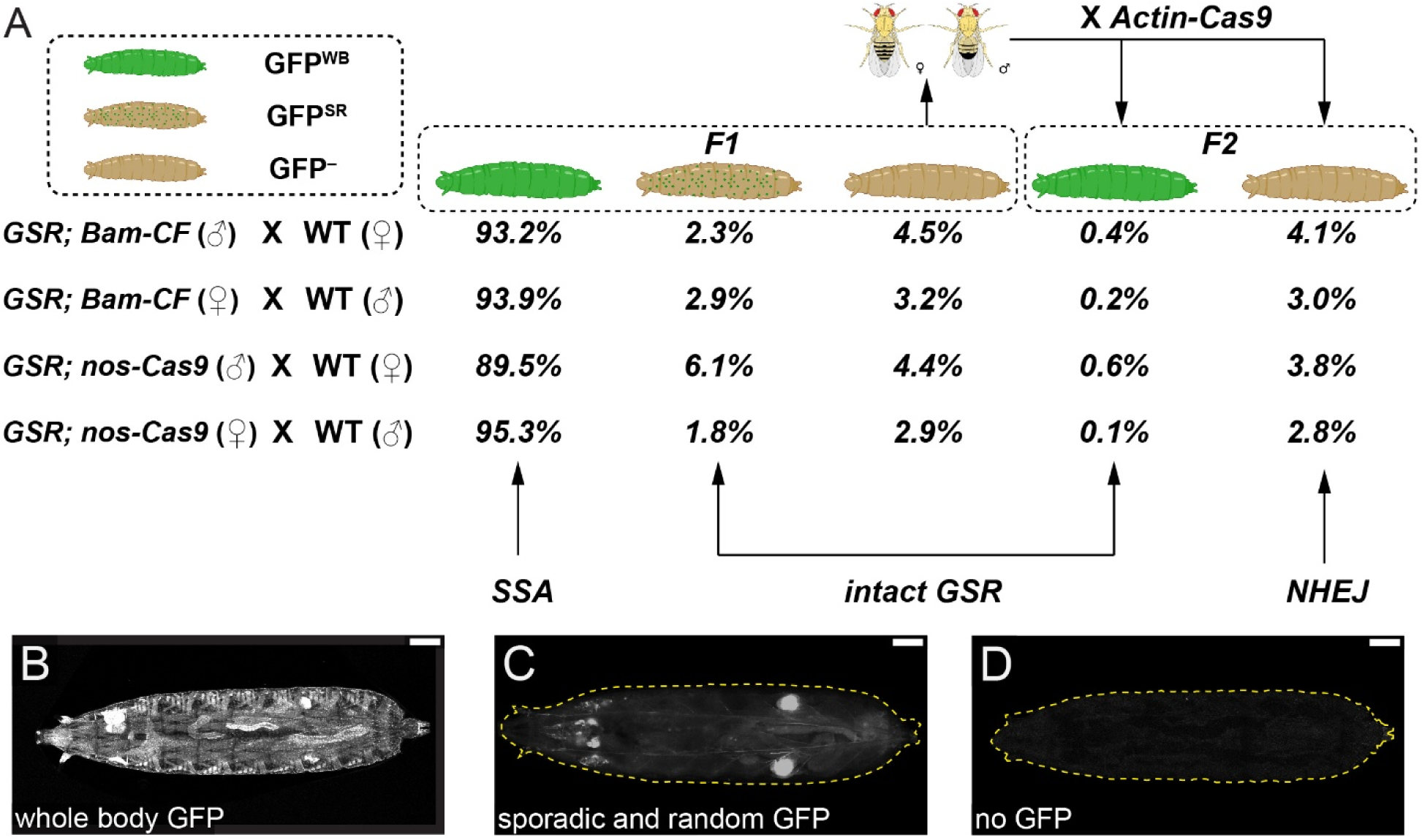
Germline Cas9 activities and DNA repair preference. (A) Diagram and data of the germline Cas9 activities tests. n = number of larvae containing *GSR*. *GSR; Bam-CF* (♂): (n = 2456), *GSR; Bam-CF* (♀): (n = 1778), *GSR; nos-Cas9* (♂): (n = 2548), *GSR; nos-Cas9* (♀): (n = 2793). (B and D) Representative images of larvae with whole-body GFP (B), sporadic and random GFP (C), and no GFP (D). For all whole larvae images, scale bar, 500 µm.

**Figure S2.**
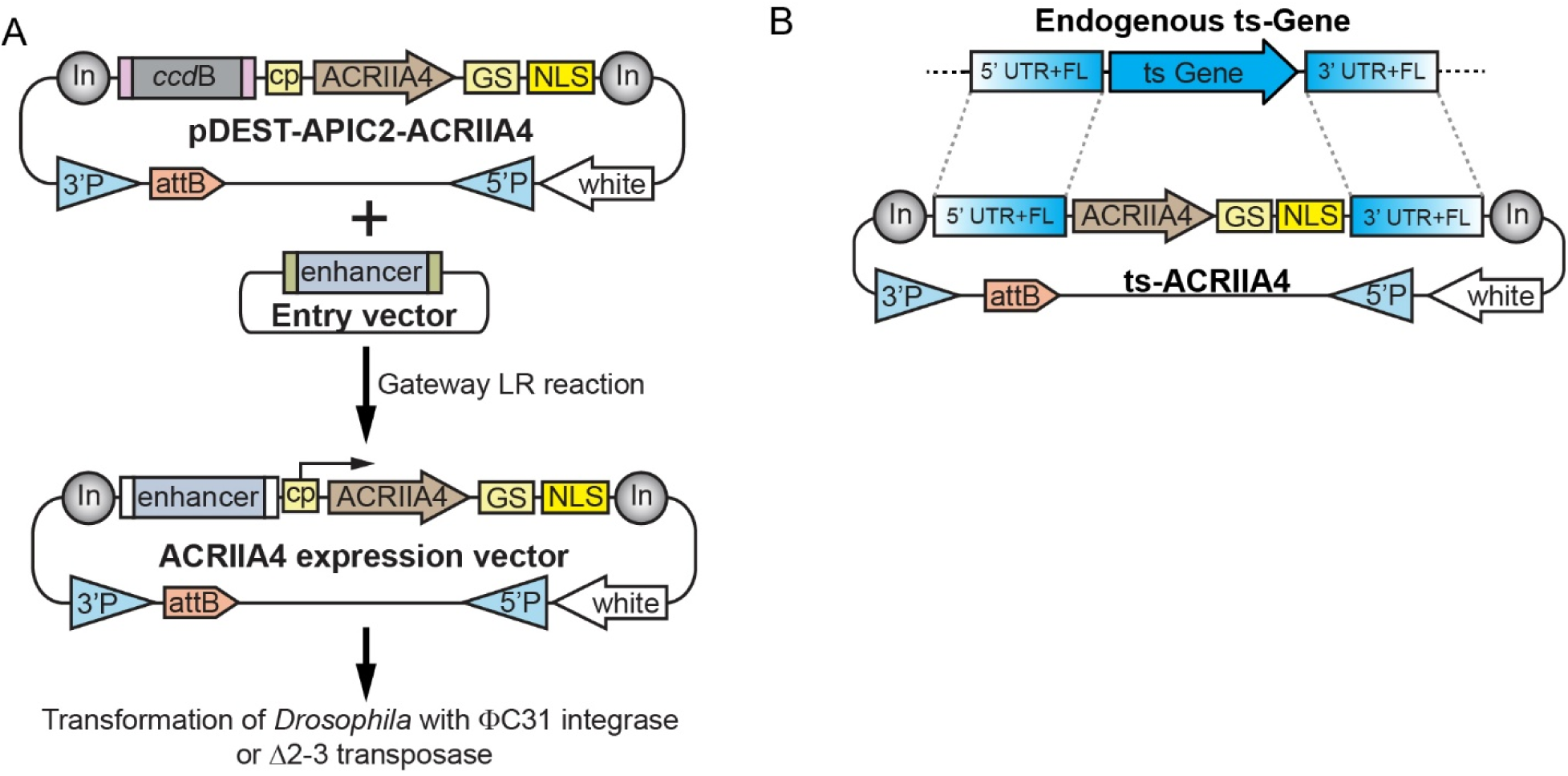
Diagram of generating tissue specific AcrIIA4. (A) Diagram of Gateway cloning and transgenesis of AcrIIA4 expression vectors. In, Gypsy insulator; cp, core promoter; 3′P and 5′P, P-element sequences; GS, glycine and serine linker; NLS, Nuclear Localization Signal. (B) Diagram of how *RabX4-AcrIIA4* and *nos-AcrIIA4* were made. Ts, tissue-specific; FL, flanking sequence.

## Notes

### Competing Interest Statement

The authors have declared no competing interest.

